# Parsing contributions of physical phenomena to smFRET statistical inhomogeneity via multiparameter stochastic simulations

**DOI:** 10.64898/2025.12.09.692639

**Authors:** Aiyan Brown, Claudiu C. Gradinaru

## Abstract

Single-molecule Förster Resonance Energy Transfer (smFRET) affords access to nanometre-scale structural and kinetic information for individual biomolecular species. Conventional analyses presuppose a strict separation of the underlying dynamical processes into distinct timescales - an assumption that is frequently violated and seldom verifiable *a posteriori*. To address this limitation, we present an integrated Brownian dynamics/stochastic simulation framework that treats the three principal dynamic contributors to smFRET observables - (i) diffusion of the molecule’s centre of mass, (ii) photophysical state-cycling, and (iii) intramolecular diffusion - in a fully time-resolved manner. Each contribution can be selectively activated, deactivated, and parametrically adjusted, thereby providing a controlled computational testbed for determining the extent to which distinct dynamical contributions alter smFRET data. By systematically varying these contributions, the individual and collective impact of specific physical processes on smFRET measurements can be delineated, and therefore the biologically-relevant information (iii) can be more precisely estimated.

## I. INTRODUCTION

Experimental molecular biophysics seeks to answer questions regarding the local and global structure of biomolecules. A common strategy involves the chemical attachment of two spatially distinct fluorescent molecules, *fluorophores*, to these molecules-of-interest. When one of the fluorophores, the *donor*, is in an electronic excited state, it may undergo a distance-dependent, non-radiative energy transfer to the nearby *acceptor* fluorophore, a process known as Förster Resonance Energy Transfer (FRET) [1].

To allow for the extraction of information beyond the ensemble average, single-molecule FRET (smFRET) techniques reduce the sample concentration to sub-nanomolar levels such that fluorescence contributions due to individual biomolecules can be individually resolved. The two most common modalities for smFRET consist of total internal reflection (TIRF) and confocal excitation schemes [2, 3]. In the latter, which will be the focus of this paper, the excitation laser beams are tightly focused into a femtolitre-scale confocal volume. Fluorescently-labelled biomolecules within the dilute sample then diffuse stochastically through this volume, producing discrete “bursts” of fluorescence: brief intervals of high photon emission rates while a labeled molecule resides in the focus, separated by intervals of background-level intensity when the molecule has diffused away.

The distance-dependent nature of FRET suggests the possibility for an inversion of the data; ratios of photons emitted by donor and acceptor fluorophores, respectively, being used as estimators of the distance between them.

However, biomolecules exist at a finite temperature and hence are generically dynamic in nature, which complicates this procedure. Experimental considerations further enforce the non-bijectivity of this problem due to, amongst other factors, finite photon budgets and instrumental temporal resolutions.

In light of this, accurate computational approaches prove to be a useful tool for interpreting data, benchmarking analysis pipelines, and disentangling the contributions of distinct physical processes to the observed signals. Current computational approaches for these sm-FRET simulations can broadly be classified into two categories. The first – and by far the most common – are coarse-grained, Monte-Carlo-based simulators operating at microsecond resolution, in which one or more molecules diffuse within a finite volume, bounded by reflective or absorptive walls [4–7]. By design, these schemes average over any dynamics occurring faster than the integration time step; consequently, nanosecond-scale processes such as laser pulse structure, rapid polymer reorientations, and fluorophore-linker motions are effectively integrated out. However, such ultra-fast motions modulate the instantaneous FRET efficiency, introducing a nontrivial convolution with slower conformational dynamics. In many cases, therefore, there is no compelling *a priori* justification for neglecting these degrees of freedom.

The second category comprises molecular dynamics simulations [8**–**10] which resolve all relevant highfrequency motions explicitly. While conceptually attractive, simulating the thousands of millisecond-long observation windows typical of confocal smFRET experiments remains far beyond current computational capabilities. Moreover, the selective activation, deactivation, or parametric adjustment of individual dynamical components is itself computationally prohibitive, often limiting the ability of such simulations to guide experiments – instead relegating them to the role of post hoc interpretation.

Bridging the gap between atomistic detail and coarsegrained efficiency is thus important for quantifying how dynamical processes shape experimentally accessible observables. We introduce here an integrated Brownian dynamics/stochastic-kinetics framework for smFRET simulation, which feasibly allows for the propagation of molecular translational and orientational degrees of freedom on sub-nanosecond time steps. The resultant trajectories feed a continuous-time, single-molecule-optimized Gillespie algorithm [11] that governs photon absorption, emission, and non-radiative transitions. By coupling Brownian dynamics with stochastic photophysics, the framework allows for one to retain the fine temporal structure necessitated by high-frequency motions, yet remains computationally tractable for simulating the tens of seconds of experimental time required to accumulate statistically meaningful data sets.

The first half of this article, Sec. II, develops the theoretical foundations of the simulation framework, developed in the Julia programming language [12], and summarizes key implementation details. The remainder is dedicated to examining how, under pulse-interleaved excitation (PIE) [13] – one of the most common confocal smFRET implementations – photophysical triplet shelving of the fluorophores and conformational dynamics of the biomolecules, both in the continuum and discrete-state limits, imprint on smFRET observables. Throughout, we connect model parameters to experimental readouts, highlight regimes of parameter identifiability and ambiguity, and provide practical guidance for interpreting smFRET data in the presence of coupled photophysical and conformational dynamics.

## II. THE SIMULATION FRAMEWORK

As first described by Förster, the instantaneous rate of donor-to-acceptor energy transfer scales as an inverse sixth power of the donor-acceptor separation *R*(*t*) := ||***R***(*t*)|| [14],

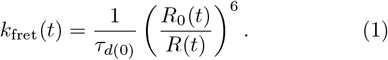

Here, the lifetime *τ*_*d*(0)_ denotes the donor excited-state lifetime in the absence of an acceptor and *R*_0_ the Förster radius, which depends upon time through the orientational factor *κ*,

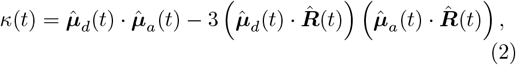

for donor and acceptor transition dipole unit vectors 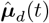 and 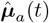. In the isotropic limit, whereby one assumes the dipoles rotate uniformly and on timescales much shorter than other sources of dynamics [10], one may approximate that the orientation is fixed, 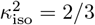.

The lifting of this assumption in the context of Brownian dynamics is discussed in some detail in Appendix A.

Our goal is hence to develop effective, dynamical models for ***R***(*t*). To do so, we must also account for the excitation mechanism that prepares the fluorophores for FRET. In the weak-excitation, on-resonance limit of the steady-state optical Bloch equations [15], a monochromatic field of angular frequency *ω* and polarization 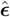 producing local intensity *I* (***X***, *t*) at position ***X*** = (*x, y, z*)^*T*^, yields an excitation rate

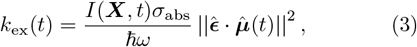

i.e., photon flux times absorption cross section, modulated by the polarization-dipole alignment. In the aforementioned isotropic limit of the dipole, the polarization-dipole alignment contributes a factor of 1*/*3, 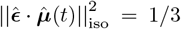, whereas slower translational diffusion of the emitter position ***X***(*t*) through the confocal point-spread function produces the characteristic burst structure via the radially decaying intensity profile.

Neglecting motions of the linker and fluorophore at present, the principal dynamic components contributing to smFRET measurements are (i) centre of mass motion of the molecule, which governs ***X***; (ii) internal translational motions that modulate the donor-acceptor separation ***R***; and (iii) photophysical phenomena altering the capacity for fluorescence emission and energy transfer processes. In what follows, we formulate physically grounded models for each component and show how they map to the rate expressions in Eqs. (1) and (3), and ultimately to the experimentally accessible observable, the photon time series.

### A. Centre-of-mass motion

In a confocal microscope, a fluorescence burst arises when a labelled molecule diffuses into the confocal volume, dwells briefly, and diffuses back out. To mimic this behaviour numerically without simulating an unbounded domain, we condition a Brownian motion on starting at a given position outside the confocal volume, diffusing to a position within it, and subsequently diffusing back outside.

To start, we consider one Brownian motion occurring over time *T >* 0 with start and end positions ***X***_0_ and ***X***_*f*_, respectively. As in a regular Brownian motion, the stochastic increments of each timestep remain Gaussian, in the sense of a Wiener process, but a time-dependent drift enforces arrival at ***X***_*f*_ almost surely, with the resulting process being known as a *Brownian bridge* [16]. Such a motion may be written in the form of a stochastic differential equation (SDE)

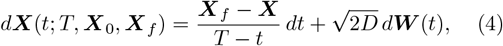

interpreted in the Itô sense, for a Wiener process ***W*** (*t*) and diffusivity *D*, subject to the initial condition ***X***(0) = ***X***_0_.

To more explicitly illustrate the simulation strategy, we define the confocal volume to be the finite region Ω_conf_ bounded by the surface whose intensity is 1*/e* of the maximum value, henceforth the 1*/e*-surface for brevity. We likewise understand the molecule to have left the confocal volume once it has reached the 1*/e*^5^-surface, *S*_ext_. If we are to sample two elements from this surface, ***X***_0_, ***X***_*f*_ ∈ *S*_ext_, and an element from the confocal volume, ***X***_1_ ∈ Ω_conf_, concatenating the two bridges ***X***(*t*; *T*_1_, ***X***_0_, ***X***_*i*_) and ***X***(*t*; *T*_2_, ***X***_*i*_, ***X***_*f*_ ), reproduces our aforementioned phenomenological picture of a burst, as shown in Fig. 1a. For all simulations performed here, the excitation beam was assumed to be a three-dimensional Gaussian with diagonal covariance matrix Σ = diag(*R*^2^, *R*^2^, (*sR*)^2^), where 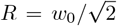 is the 1*/e* beam width and *s* the structure factor. As a result, surfaces of constant intensity are prolate spheroids.

**FIG. 1.**
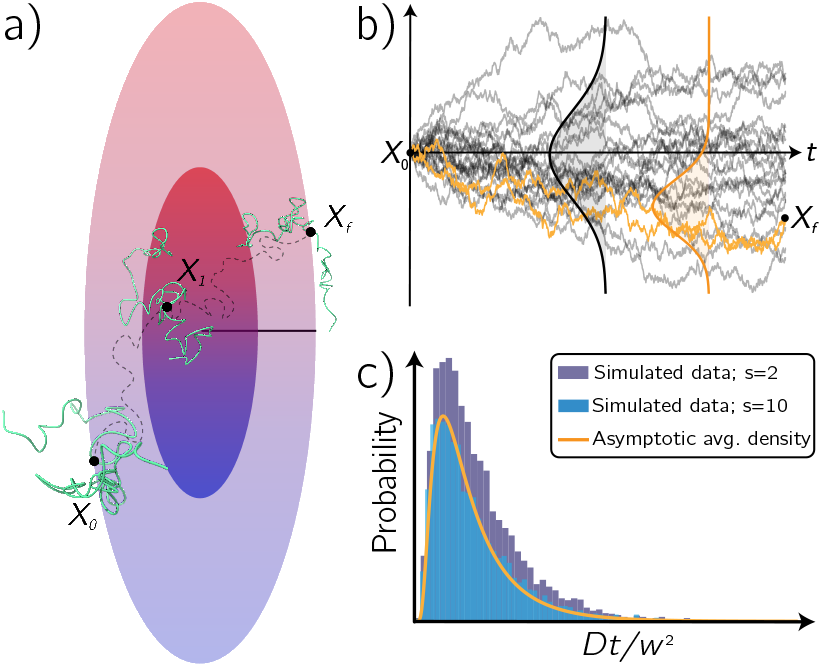
Outline of Brownian bridges and their usage as the COM motion of the biomolecule. (a) Sample COM trajectory starting outside the confocal volume at X0, traversing inside to X1, and traversing outside once more to Xf . Molecular graphics in this figure and all henceforth were performed with UCSF ChimeraX, developed by the Resource for Biocomputing, Visualization, and Informatics at the University of California, San Francisco [17]. (b) Brownian bridge trajectories (orange) compared to un-conditioned Brownian motions (grey) and snapshots of the time-dependent probability densities (vertical curves). (c) Comparison of the asymptotic 3D Bessel process exit-time distribution (orange) Eq. (5) and a Monte-Carlo simulated exit time distribution from a 2:1 (purple) and 10:1 (blue) prolate sphereoid.

To determine the time allocated to each bridge, *T*_1_ and *T*_2_ above, we define the exit time of a Brownian motion ***X*** from a bounded volume 𝒱 starting at a position ***X***_0_ ∈ 𝒱 as 𝒯_*𝒱*_ (***X***_0_) := inf {*t* ≥0 : ***X***(*t*) ∈*∂* 𝒱 | ***X***(0) = ***X***_0_ }. Sampling from the distribution of exit times from the region bounded by the 1*/e*^5^-surface, starting at ***X*** ∈ Ω_conf_, should hence preserve the diffusion statistics of the bursts. Since no closed form for the exit-time distribution from a three dimensional ellipsoid is known, as far as the authors are aware, we approximate by sampling each time from a rescaled (asymptotic) exit time distribution of a three-dimensional Bessel process. This choice can be applied in both the forward and reverse cases due to the time-reversal symmetry of the underlying Brownian motion.

It is convenient to perform an affine rescaling of position variables, 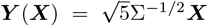, where the factor of 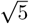 accounts for the fact that we are exiting from the 1*/e*^5^-surface. Starting at the radial distance *y* = ||***Y*** (***X***_1_) ||, Brownian bridge durations were sampled from [18]

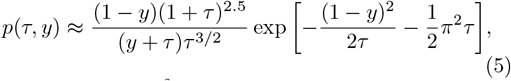

where 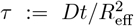 is the non-dimensionalized time. The effective radius 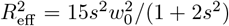, the scaled harmonic mean of the squared semi-axes of the ellipsoid, was chosen to preserve the mean first-passage time, 𝔼 [𝒯 _*𝒱*_ (***X***_0_)], of the exit-time distribution under the change of variables, as derived in the supplementary material.

To validate this approximation for the exit time, Monte Carlo simulations were performed for various spheroid aspect ratios, Fig. 1c. We observe that the distribution of Eq. (5) asymptotically approximates the genuine exit time density for increasingly prolate excitation profiles.

### B. Translational dynamics

At thermodynamic equilibrium, without loss of generality, the time evolution of the donor-to-acceptor fluorophore displacement ***R*** is determined by a Langevin equation in a potential energy landscape *U* . We assume that these motions are overdamped, such that the contribution due to inertia is negligible, and the diffusivity *D* is homogeneous such that we may write ***R***(*t*) as an SDE

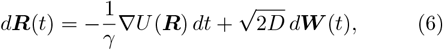

interpreted in the Itô sense, for damping coefficient *γ*, related to diffusivity by a fluctuation-dissipation theorem (FDT) *Dγ* = *k*_*B*_*T* .

In the isotropic limits of Eqs. (2) and (3), only distance fluctuations are relevant to the energy transfer rate Eq. (1). In light of this, we likewise assume that the potential *U* is rotationally isotropic such that it may treated as a one-dimensional function in the separation distance *R*(*t*). By Itô’s lemma [16], the differential equation for *R* now reads

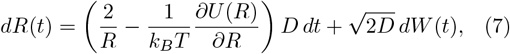

including a repulsive drift 2*D/R*, the Itô correction, which enforces positive support of *R*.

Suppose that *U* (*R*) observes a local minima at some radius *R*_eq_. If we set *U* (*R*_eq_) = 0, then we may locally approximate the potential as a quadratic in (*R* − *R*_eq_)^2^. Upon substitution into Eq. (7), we get the equation for a modified Ornstein-Uhlenbeck (OU) processes, i.e., a “stochastic spring”,

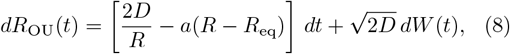

where we have defined the rate of mean reversion *a* = ∇ ^2^*U* (*R*_eq_)*/γ*. In the particular case where we set *R*_eq_ = 0, the equilibrium distribution reduces to the classical freely-jointed-chain form. Eq. (8) acts as an effective approximation for the behaviour of a system with a minimal structure and a single, perhaps poorly defined, minima, such as is in the case of a single-stranded DNA or a disordered protein.

When *U* exhibits multiple minima, the dynamics depend upon the global curvature of the landscape, which is more challenging to model in general. We hence break from the continuum picture into discrete states, assuming negligible fluctuations about the minima in this case, and Markovian discrete-state description with a continuous-time transition-rate matrix governing conformational switching among the finitely many states. This approach has been widely used in smFRET analyses to date (c.f., Refs. [4, 7]), and will be discussed in detail in the next subsection.

### C. Determining photon emissions

With ***X***(*t*) known from the COM dynamics of Sec. II A, we model the excitation field as a pulsed train modulated by the confocal point-spread function (PSF) *G*. For a repetition rate *ν* and pulse envelop *p*(*t*), the local intensity of a given excitation source is

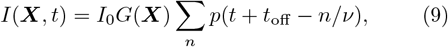

where *I*_0_ sets the peak power and *p* is the temporal pulse shape. As alluded to in Sec. II A, we model the confocal PSF by a separable three-dimensional Gaussian distribution,

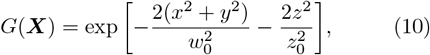

with lateral beam waist *w*_0_ and axial parameter *z*_0_ = *sw*_0_. We furthermore take *p*(*t*) to be Gaussian with full width half max *τ*_*p*_ ≪ 1*/ν*; for pulse interleaved excitation (PIE), to achieve a 1:1 duty cycle, the temporal offset is set to 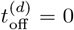 and 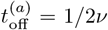 for donorand acceptorselective pulses, respectively, as in Fig. 2c.

**FIG. 2.**
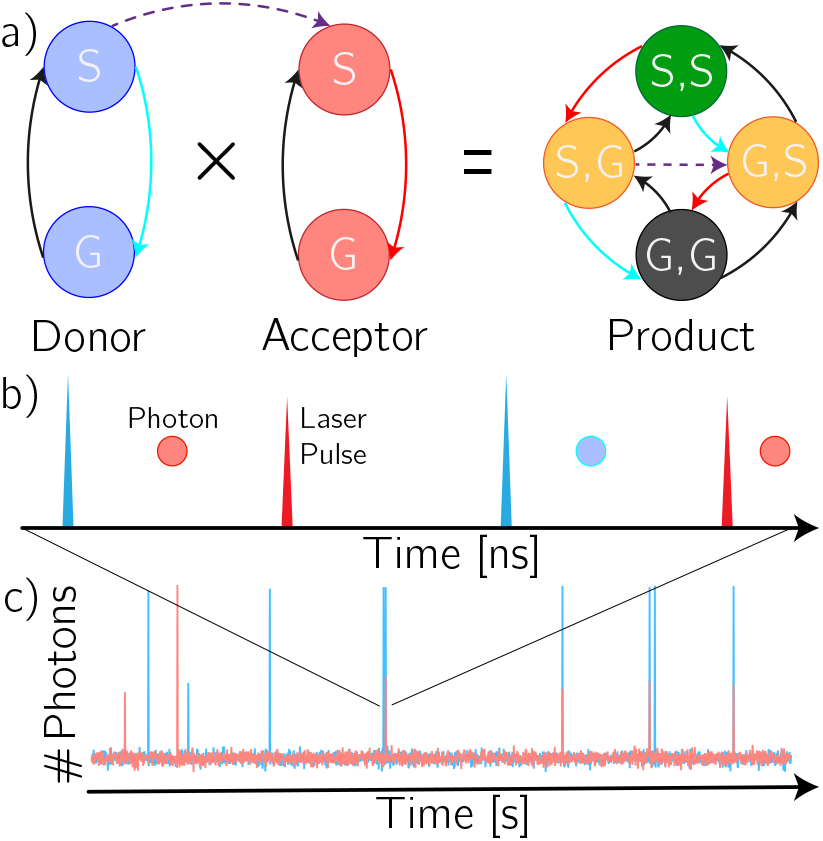
Example of the procedure for generating smFRET data. (a) Two minimal multidigraphs – for the donor and acceptor, respectively – and their four-node (combined) product graph. Blue and red edges denote fluorescent emission pathways, while the orange/ dashed edges represents the FRET pathway. (b) Generation of photons, via de-excitation along fluorescent pathways in (a), due to pulsed laser excitation. (c) Long-time confocal smFRET data structure: background noise amongst high photon rate “bursts”.

The instantaneous FRET and the two excitation rates, Eqs. (1), (3), are three of many possible photophysical channels which a pair of fluorophores may react by. To see how this complexity arises, we first consider the case of a single fluorophore with a set *V* of electronic states. We formalize the photophysical reaction network as a labelled multidigraph,

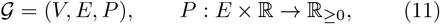

where *E* ⊂*V* ×*V* a multiset of directed transitions (edges), and *P* (*e, t*) assigns to every edge *e* = (*i, j*) its instantaneous rate or *propensity* at time *t*. For instance, a single fluorophore with a ground and a singlet state, *G* and *S* respectively, can be written *V* = {*G, S}* and *E* = (*G, S*), (*S, G*) with *p*((*G, S*), *t*) = *k*_ex_(*t*) and constant fluorescence rate *P* ((*S, G*), *t*) = *k*_fl_ describing transitions to *S* from *G* (excitation) and vice versa (de-excitation). In the simulation of molecular systems with discretely many states, the propensities are assumed time-independent, and a single reaction network 𝒢 sufficient to capture the dynamics.

The joint network of two non-interacting fluorophores 𝒢_1_ = (*V*_1_, *E*_1_, *p*_1_) and 𝒢_2_ = (*V*_2_, *E*_2_, *p*_2_) is obtained by the product 𝒢_1_×𝒢_2_ = (*V*_1_ × *V*_2_, *E*_1_ *E*_2_, *P*_1_ *P*_2_), where an edge ((*i*_1_, *i*_2_), (*j*_1_, *j*_2_)) ∈*E*_1_ *E*_2_ exists if either (*i*_1_, *j*_1_) ∈*E*_1_ with *i*_2_ = *j*_2_ or (*i*_2_, *j*_2_) ∈*E*_2_ with *i*_1_ = *j*_1_. If we take the first index to denote the donor state and the second the acceptor, energy transfer is introduced by adding an edge only defined on the joint network, *e*_fret_ = ((*S, G*), (*G, S*)) and *P* (*e*_fret_, *t*) = *k*_fret_(*t*), which transforms an excited donor into an excited acceptor state with rate *k*_fret_, see Fig. 2. This construction preserves the tensor-product topology of independent processes while superimposing the multi-fluorophore couplings required to reproduce the full electronic-state kinetics of smFRET experiments.

For a given photophysical network 𝒢 = (*V, E, P* ), let the system reside in the state *X* ∈ *V* at time *t*. We define the source and destination maps *s* : *E* → *V*, (*v*_1_, *v*_2_) → *v*_1_ and *f* : *E* → *V*, (*v*_1_, *v*_2_) → *v*_2_ respectively, and denote by

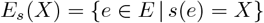

the set of admissible transitions from *X*. Suppose that each outgoing edge *e* ∈ *E*_*s*_(*X*) is treated as a, potentially time-inhomogeneous, Poisson process with instantaneous propensity *P* (*e, t*). Following the work of Anderson [19], the cumulative distribution function for the waiting time Δ to the next reaction is therefore

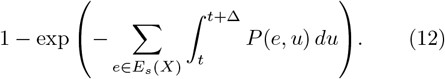

Per-transition waiting times Δ_*k*_, for each *k* ∈ ℕ indexing the set *E*_*s*_(*X*), are generated by inverse-transform sampling Eq. (12),

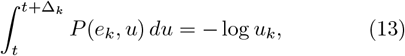

with *e*_*k*_ ∈ *E*_*s*_(*X*) and for uniform random variables *u*_*k*_ ∼ 𝒰 (0, 1). The imminent transition/reaction is the edge with the smallest value for Δ_*k*_; *m* = arg min {Δ_*k*_} The simulation time and state are updated, *t* → *t* + Δ_*m*_ and *X* → *f* (*e*_*m*_), and the above algorithm is repeated until: (a) the time allocated has been reached or exceeded, or (b) a sink is reached, i.e., a node with outdegree zero. Eq. (13) is integrated numerically unless *P* (*e*_*k*_, *t*) = *P* (*e*_*k*_) is time-independent, in which case the closed-form expression Δ_*k*_ = − log *u*_*k*_*/P* (*e*_*k*_) follows trivially.

## III. THE EFFECT OF TRIPLET STATES

A minimal description of fluorophore photophysics treats the system as a two-state model in which the molecule resides either in the ground state *G* or an excited singlet state *S*, populated by photon absorption, as in Fig. 2a. From *S*, the system returns to *G* by radiative decay at a rate *k*_fl_ = *φ*_fl_*/τ*_fl_ or by internal conversion at rate *k*_ic_ = (1−*φ*_fl_)*/τ*_fl_, where *φ*_fl_ and *τ*_fl_ are the fluorescence quantum yield and lifetime, respectively. While convenient, this scheme omits long-lived “dark states” that transiently or permanently suppress fluorescence – most prominently in the form of a triplet state *T* and an irreversibly photobleached state *B*.

Photobleaching is effectively captured by extending the two-state graph to include *T* and a photoexcited triplet manifold *T* ^*^ that mediates access to *B* (cf. Ref. [20]). In a standard five-state picture, excitation promotes *G* → *S* at the time-dependent rate *k*_ex_(*t*), Eq. (3), *S* decays radiatively, *k*_fl_, or non-radiatively, *k*_ic_, and intersystem crossing (ISC) populates *T* at a rate *k*_isc_ = *φ*_isc_*/τ*_fl_, with an ISC quantum yield *φ*_isc_. The triplet state relaxes thermally to *G* with a rate equal to the inverse of its lifetime *τ*_tr_, but may also absorb another photon with effective cross section *σ*_tr_, promoting *T* → *T* ^*^, from which an irreversible channel to *B* operates with *k*_bl_. Thus the ef-fective bleaching hazard increases with the instantaneous intensity *I*(***X***, *t*), generating a practical trade-off: higher excitation yields more photons per unit time but shortens the observation window.

The triplet lifetime *τ*_tr_ ∼ 10^−5^ s typical of fluorophores used in smFRET measurements is comparable to many biomolecular timescales, so triplet blinking itself can imprint kinetics on intensity traces and on dwell-time statistics. Recent work by Pati *et al*. [21] showed that triplet occupancy in either donor or acceptor fluorophores biases FRET efficiencies in TIRF-based smFRET. As a first illustration of our burst-simulation framework, we adopt the analogous photophysical scheme, whereby both fluorophores photophysics follow a three-state {*G, S, T* } subgraph, as shown in Fig. 3a. Conditional on not being in *T*, the addressed dye is excited at the appropriate *k*_ex_(*t*), while FRET proceeds at a constant rate 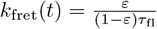 for input FRET efficiency *ε*.

**FIG. 3.**
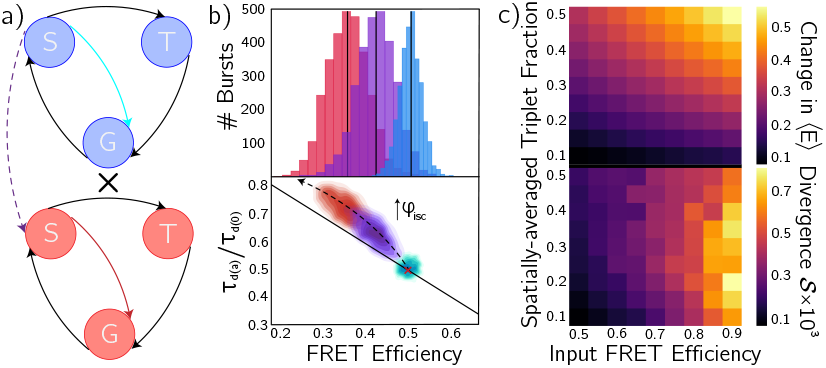
Characterization of the effect of acceptor triplet states on observables in smFRET with PIE with fixed triplet lifetime *τ*_tr_ = 10 *µ*s. (a) Minimal model of triplet state photophysics, consisting of a ground (*G*), singlet (*S*) and triplet (*T* ) state for each fluorophore. (b) Normalized apparent donor lifetime in the presence of acceptor versus FRET efficiency (bottom) and its marginal (top) distribution. In both cases, a shift in the FRET efficiency and broadening of the distributions is observed when the acceptor intersystem crossing quantum yield, *φ*_isc_, increases. (c) Summary of the effect of acceptor triplet states on the first (top) and higher-order (bottom) moments of the FRET distribution.

In the PIE scheme, the estimator for FRET efficiency from photon counts, the proximity ratio, is

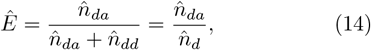

where 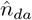 (resp. 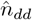) counts acceptor-channel (resp. donor-channel) photons following donor-targeted pulses.

Because the presence of a donor triplet state suppresses both channels during the donor-excitation windows for which it is occupied, it cancels in the ratio Eq. (14) and does not bias *Ê*. By contrast, the acceptor triplet state disables the capacity for transfer, while increasing donor radiative decay, thereby downshifting *Ê* and skewing FRET histograms towards lower efficiencies, Fig. 3b.

If the acceptor triplet state is occupied a fraction 0 *< f*_tr_ *<* 1 of the time the donor is excited, the expected shift in the mean FRET efficiency is expected to be linear in both the fraction and input FRET efficiency, − *f*_tr_*ε*, compensating for the mean shifts 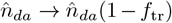 and 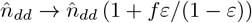 – see the Supplementary material for a detailed derivation. This calculation is complicated by the explicit dependency of the triplet occupation fraction on the excitation rate, and hence position and time:

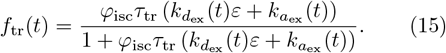

The mean intensity conditioned on detection,

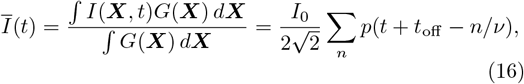

was used in place of *I*(***X***, *t*) in Eq. (3), resulting in a decoupling of the excitation rates from the trajectory. Subsequently performing a time-averaging of Eq. (15), one can determine the mean triplet fraction 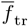, which we used to determine the ISC quantum yields to perform simulations at.

Simulations were performed for various values of the independent variables determining triplet state behaviour, namely the triplet fractions 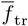 and the input FRET efficiencies *ε*, with no additional sources of input dynamics – these will be discussed in the following section. As expected, for fixed input FRET efficiency or triplet fraction, a monotonic and near linear – see Supplementary material for analysis of the marginal distributions – increase in the change in the mean efficiency is observed, amounting to systems with higher *a priori* FRET efficiencies being more significantly influenced by the presence of acceptor triplet states, Fig. 3c. In effect, the product of the photophysical networks and the single-node conformational network amounts to a two-state continuous-time Markov chain (CTMC) with *E* = *ε* and *E* = 0 nodes with forward and reverse rates 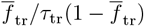 and 1*/τ*_tr_, respectively (see Sec. IV A).

More complex are the beyond-first-order changes in the FRET efficiency. We quantify these changes by calculating a per-photon Kullback-Leibler divergence, = 𝒮 ⟨*N*⟩ ^−1^*D*_KL_ (*P* −*µ*_*P*_ ||*Q* −*µ*_*Q*_), between the simulated *P* and shot-noise-limited *Q* distributions, calculated follow Nir *et al*. (2006) [4], for an average number of photons per burst ⟨*N*⟩ . Larger 𝒮 indicates more significant differentiation between the two distributions – a more significant contribution from dynamics to the FRET distribution. As discussed in the supplementary material, we understand the near-monotonic relationship between divergence and FRET efficiency to be a consequence of the variance, while the non-linear behaviour of 𝒮 as a function of triplet fraction shown in Fig. 3c implies significant contributions from higher-order moments.

## IV. MOLECULAR DYNAMICS

Photophysics, though interesting and highly-nontrivial in its own right, are understood as a secondary player in smFRET measurements. The dynamics of interest are due to conformational changes in the molecule itself, but it is not well understood the extent to which the presence of triplet states – inducing their own apparent dynamics in the sense of the previous section – convolute these measurements. We begin by considering the well-studied case whereby conformational dynamics are parameterized by a continuous-time Markov chain, with nodes mapping to fixed FRET efficiencies, and investigating the effect of acceptor dark states across disparate conformational timescales. This informs our study of continuous conformational dynamics, modelled via a non-linear stochastic spring. First, the effects of the free energy landscape and diffusivity are differentiated in the absence of the triplet state. Subsequently, the triplet state is introduced and the changes of the distributions are characterized relative to the characteristic timescale of the stochastic process governing the conformational dynamics.

### A. Discrete-state systems

Consider a molecule whose conformational dynamics can be effectively described by exchange between two discrete states with FRET efficiencies *E*_1_ and *E*_2_, interconverting with forward and backward rates *k*_*f*_ and *k*_*b*_, respectively. As discussed previously (cf. Refs. [7, 22–24]), the observed FRET histograms depend on both the ratio *k*_*f*_ */k*_*b*_, which sets the equilibrium occupancy of the two states, and the total exchange rate 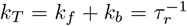, which defines the characteristic interconversion time *τ*_*r*_.

In the limit *τ*_*r*_ → 0, where conformational exchange is much comparable to the mean inter-photon time, the measured FRET efficiency approaches the population-weighted average of *E*_1_ and *E*_2_. In this regime, the presence of dynamics is manifested by a distribution whose breadth asymptotically approaches that of the shot-noise limit (SNL), together with an upward deviation from the static FRET line in a lifetime-versus-FRET plot. As *τ*_*r*_ increases, exceeding to the inter-photon time but remaining less than the burst duration, the distribution broadens further and appears to interpolate between *E*_1_ and *E*_2_ along the dynamic FRET line [22]. In the opposite limit of slow exchange, where *τ*_*r*_ approaches the typical burst duration, two nearly shot-noise-limited peaks emerge at *E*_1_ and *E*_2_ (see Fig. 4b). We refer to these three limits as the fast, intermediate, and slow dynamic regimes of smFRET, respectively.

**FIG. 4.**
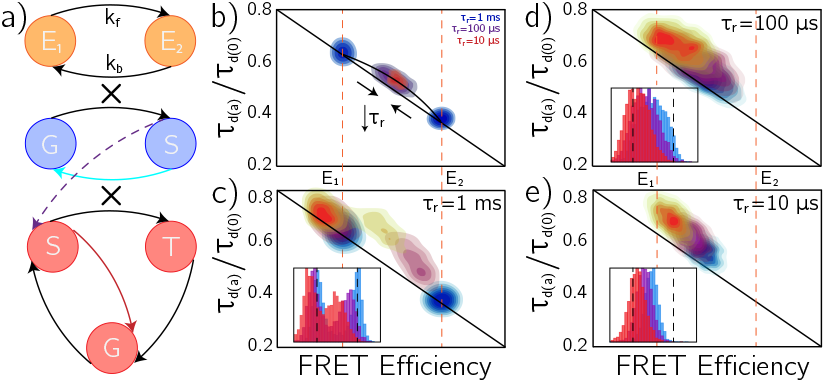
Effect of an acceptor triplet state on apparent twostate conformational dynamics as reported by confocal smFRET. (a) Schematic of the coupled conformational and photophysical state manifolds. (b) Normalized donor lifetime versus FRET efficiency for various interconversion lifetimes *τ*_*r*_ in the absence of triplet blinking. As *τ*_*r*_ decreases below the mean inter-photon time, the two discrete populations merge toward their population-weighted mean along the dynamic FRET line. (c) Slow conformational-exchange regime, *τ*_*r*_ *≫ τ*_tr_. For fixed intersystem crossing quantum yield (*φ*_isc_ = 0, 1.25 *×*10^−3^, 2.75 *×*10^−3^ for blue, purple, and red/yellow curves, respectively; mean donor and acceptor excitation rates 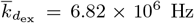 and 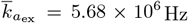), the acceptor triplet state perturbs the high-FRET population at *E*_2_ more strongly than the low-FRET population at *E*_1_. (d) Intermediate conformational-exchange regime, *τ*_*r*_ *≈ τ*_tr_. The population divergence 𝒮 decreases and the FRET histograms become increasingly homogeneous as the triplet fraction increases. (e) Fast conformational-exchange regime, *τ*_*r*_ *≪ τ*_tr_. The entire, dynamically-averaged, population is affected nearly homogeneously by the acceptor triplet state.

The triplet lifetime *τ*_tr_ typically falls within the fast-tointermediate regime, leading to a nontrivial overlap between photophysical and conformational timescales. To quantify how this interplay affects the influence of dark states, we performed simulations in which both the mean triplet fraction 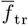, see Eq. (15), and the interconversion time *τ*_*r*_ were systematically varied. Throughout, the rate ratio *k*_*f*_ */k*_*b*_ was fixed to 1.0, so that in the absence of triplet formation the two conformations, with FRET efficiencies *E*_1_ = 0.35 and *E*_2_ = 0.65, are equally populated on average.

As briefly discussed in the previous section, the presence of an acceptor triplet state induces an effective conformational dynamics between a dark state with vanishing FRET efficiency and the underlying conformational FRET states. In an abstract sense, adding genuine twostate conformational exchange to this photophysics yields an effective four-state system, defined on the product space of conformational and photophysical states. More explicitly, suppose the conformational dynamics are described by the CTMC generator

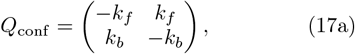

and likewise the effective photophysics by the generator

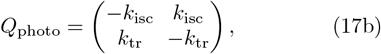

where *k*_tr_ = 1*/τ*_tr_ and 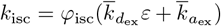, following Eq. (15). Assuming the conformational and photophysical processes are independent (i.e., no quenching or synchronized transitions), the joint dynamics on the product space are governed by the Kronecker sum of the local generators [25],

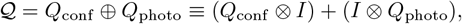

where *I* is the identity on the complementary space. Fig. 4a schematizes this four-state model, explicitly indicating the conformational and per-fluorophore photophysical transitions.

In the limit of fast photophysics, (*k*_tr_+*k*_isc_) ≫ (*k*_*f*_ +*k*_*b*_), (*k*_tr_ + *k*_isc_) ≳ *T* ^−1^ where *T* is the burst duration, the singlet-triplet sub-chain rapidly equilibrates within each conformation. The photophysical occupancy fractions are 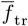 and 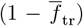, for the dark and bright states respectively, with *ε* in Eq. (15) being dictated per-conformation, amounting in an apparent triplet fraction 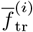 for conformation *i*. Adiabatic elimination projects onto the slow conformational manifold: to leading order, the generator *Q*_conf_ is retained, while the emissive output of conformation *i* is scaled by its bright fraction. Thus, conformation *i* appears with an apparent FRET efficiency 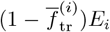. The result is two conformation-specific signatures, each diminished according to the time spent in the triplet state, as illustrated in Fig. 4c: the state sequence is dominated by slow conformational switching, producing two nearly distinct populations, while the comparatively rapid triplet blinking is averaged into a per-population attenuation of the FRET efficiency. In the opposite limit of rapid molecular reconfiguration, (*k*_*f*_ + *k*_*b*_) ≫ (*k*_tr_ + *k*_isc_), the conformational coordinate equilibrates quickly under (*Q*_conf_ ⊗ *I*) to the steady state

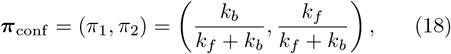

independently in each photophysical layer. A naive projection that fully lumps over conformations, in analogy with the previous limit, yields purely on/off kinetics and consequently removes the conformation-induced deviations from the static FRET line (cf. Fig. 4b). To retain these dynamic shifts while still coarse-graining, we instead lump only the two dark sub-states and keep the bright conformations distinct, leading to the three-state generator

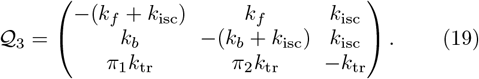

Here the return probabilities from the dark state are *π*_1_ and *π*_2_ because, under independence and rapid conformational mixing, the dark population is replenished from the bright manifold in proportion to ***π***_conf_ .

However, when the molecular reconfiguration becomes increasingly rapid, the assumption that internal dynamics are slow on the burst timescale breaks down and the finite burst duration *T* must be taken into account explicitly. As will be discussed in some detail in Sec. IV B, the analysis holding for a two-state CTMC as it does for the OU process considered in Eq. (24b), the dynamic contribution to the broadening decays as ∼ 1*/*(*k*_slow_*T* ) for *k*_slow_*T* ≫ 1 where

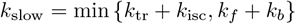

is the slowest internal relaxation rate of the joint conformational-photophysical process. The internal dynamics are therefore effectively averaged over within each burst and the observed FRET distribution approaches its SNL. In this “near-SNL” regime, the burst-averaged observables depend primarily on the stationary emission fractions from the two bright conformations and the dark pool, and the effective three-state description in Eq. (19) remains accurate even if *k*_*f*_ +*k*_*b*_ ∼ *k*_tr_ +*k*_isc_, as is the case in Fig. 4e. Here, the photokinetics remain effectively twostate (bright/dark), but the bright state carries a mixture of *E*_1_ and *E*_2_ weighted by their steady-state probabilities.

In between these cases, Fig. 4d, the separation of manifolds or asympotic behaviour underlying the coarsegrained limits breaks down, and the full four-state model must be retained to capture the dynamics generated by 𝒬. The mean FRET response in this regime remains similar to that obtained from the three-state description of rapid conformational exchange, Eq. (19), in the sense that the population-averaged efficiency still follows a dynamic FRET line between *E*_1_ and *E*_2_, and the addition of triplet induces a “third state” at *E* = 0, stemming from this dynamic line. However, the divergence of the distribution relative to the SNL decreases with increasing triplet fraction: bursts with higher apparent FRET (i.e., trajectories spending notably more time in *E*_2_ than *E*_1_) are more strongly affected by the acceptor triplet, leading to a partial homogenization of the histogram. In practice, this overlap regime is the most challenging to interpret, as photophysics and conformational exchange jointly shape the FRET statistics; if triplet blinking is not explicitly accounted for, the resulting reduction in broadening can masquerade as weaker or slower conformational dynamics than are actually present.

### B. Highly disordered systems

A molecule whose curvature about a minima in its free energy landscape is sufficiently small that it samples a broad range of FRET efficiencies can no longer be effectively described by a discrete-state model. Instead, after performing a series expansion about the minimum and introducing a nonlinear Itô drift correction to enforce positivity of the separation, the donor-acceptor distance becomes a realization of Eq. (8). The free-energy landscape corresponding to this SDE can be written as

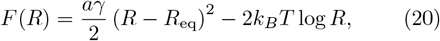

where the quadratic term represents the coarse-grained elastic confinement of the polymer and the logarithmic term arises from the radial Itô correction, which may be interpreted as an entropic contribution. Using the FDT to eliminate *γ* in favor of the internal diffusivity *D*, one finds that *F* (*R*) attains a minimum at

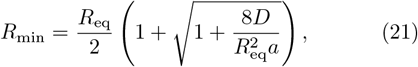

with a local effective mean-reversion rate

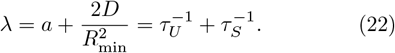

Here we have introduced the “potential” and “entropic” lifetimes *τ*_*U*_ = *a*^−1^ and 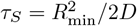, respectively. The effective reconfiguration time,

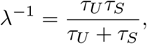

is thus the harmonic mean of these two contributions, reflecting the combined influence of energetic confinement and entropic repulsion on the relaxation of *R*.

For *R* ≈ *R*_min_, the process is approximately OU with mean-reversion rate *λ*, whose stationary covariance is

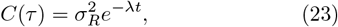

where 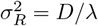 is the stationary variance of the process. The time-averaged variance over an interval of duration *T* can then be evaluated using Eq. (23) as

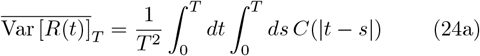

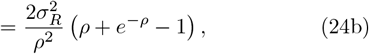

where we have defined *ρ* = *λT* as the number of effective reconfigurations occurring over time *T* . The averaging in Eq. (24a) provides a continuum analogue of the photon-by-photon averaging underlying the FRET estimate in Eq. (14) within a single burst. In this context, *T* is naturally identified with the burst duration. For *ρ* ≫ 1, the fast conformational-dynamics regime which this model typically applies to, 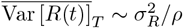 and the dynamic contribution to the variance vanishes asymptotically, reducing the observed broadening to the SNL. This behavior closely parallels the discrete-state analysis: the ability to resolve conformational timescales is dictated by the “natural” lifetimes of the experiment, namely the inter-photon time and the typical burst duration.

To assess the broadening predicted by Eq. (24b) across a range of relaxation timescales *λ*, we performed simulations in which the conformational dynamics were governed by Eq. (8). The dynamical timescales were chosen to be near those reported for intrinsically disordered proteins (cf. Ref. [26]), which can be described by canonical polymer-physics equations of motion and thus, in a coarse-grained sense, stochastically represented by Eq. (8). In the continuum simulations shown below, we varied the potential timescale *τ*_*U*_ = *a*^−1^ and internal diffusivity *D* separately, thereby modulating both the curvature of the harmonic contribution to *F* (*R*) and, through Eq. (21), the relative weight of the entropic term. This provides a controlled way to explore how energetic stiffness and entropic broadening jointly determine the effective reconfiguration time and, in turn, the observed FRET distributions.

Taken together, Fig. 5 illustrates how energetic stiffness and entropic broadening jointly control the continuum dynamics and their FRET signatures. Increasing the harmonic stiffness (decreasing *τ*_*U*_ = *a*^−1^) steepens the quadratic contribution to *F* (*R*) and pulls *R*_min_ closer to *R*_eq_, Fig. 5a, thereby suppressing the entropic term and narrowing the equilibrium distribution. At fixed *τ*_*U*_, increasing *D* enhances the weight of the entropic contribution, pushing *R*_min_ outward, Eq. (21), and broadening the accessible range of separations, as seen directly in the time traces of Fig. 5b. Panels c-d show how these microscopic changes map onto the observables: larger *D* at fixed *τ*_*U*_ (c) or larger *τ*_*U*_ at fixed *D* (d) both increase the effective reconfiguration time *λ*^−1^ and the amplitude of distance fluctuations on the burst timescale, leading to broader FRET distributions, reduced mean FRET efficiencies, and a more pronounced dynamic shift above the static FRET line.

**FIG. 5.**
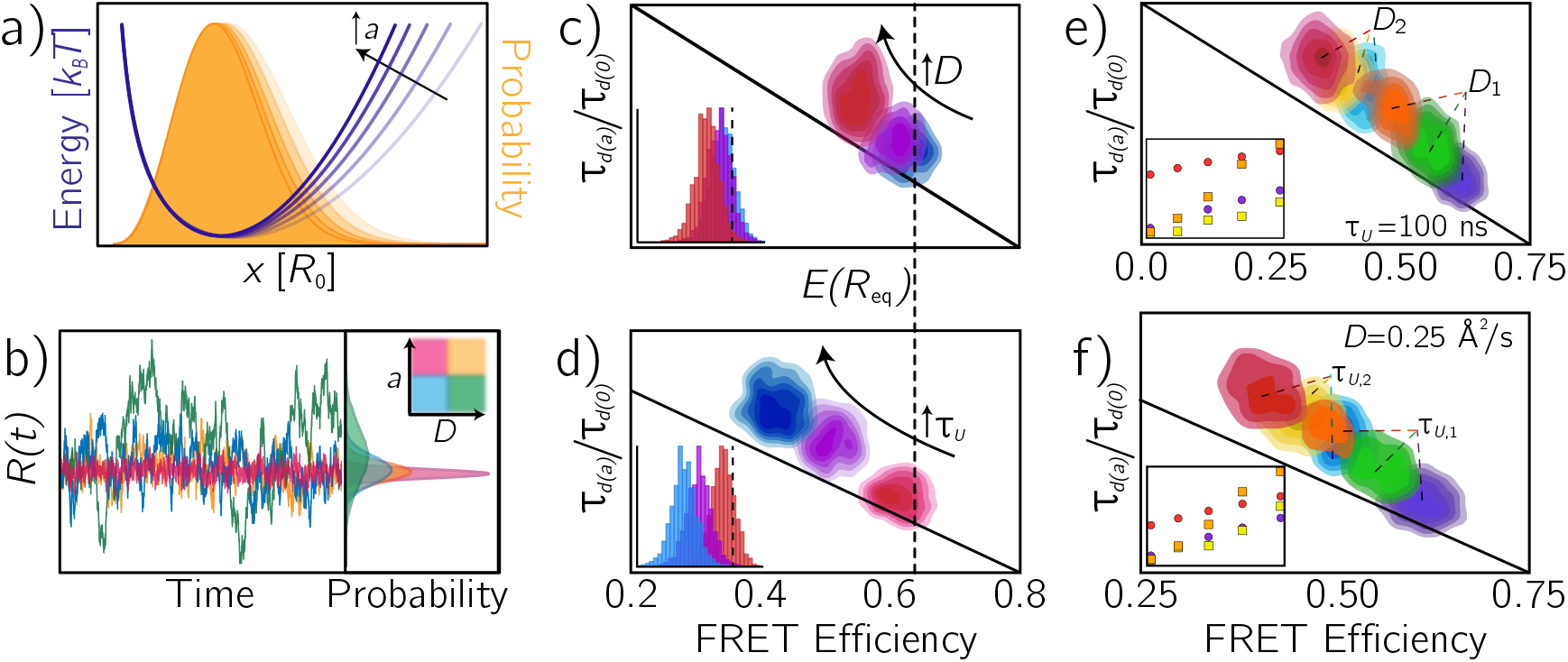
Stochastic continuum dynamics of a highly disordered molecule in the absence and presence of an acceptor triplet state. (a) Effect of varying the mean-reversion rate *a* on the effective free-energy landscape *F* (*R*) (blue) and the corresponding equilibrium probability density (orange). Larger *a* leads to a steeper well and a narrower equilibrium distribution of *R*. (b) Representative time traces of the donor-acceptor separation *R*(*t*), Eq. (8), for different combinations of internal diffusivity *D* and mean-reversion rate *a*, illustrating the approach to the local minimum and the fluctuation amplitude about it. (c) Normalized donor fluorescence lifetime versus FRET efficiency histograms for varying internal diffusivity in the absence of triplet blinking. As *D* increases (10 Å^2^ */*ns, red; 6.25 Å^2^ */*ns, purple; 1.0 Å^2^ */*ns, blue) at fixed potential timescale *τ*_*U*_ = 10 ns, the FRET distribution broadens, the mean FRET efficiency decreases, and the dynamic shift above the static FRET line becomes more pronounced. Inset: corresponding FRET-efficiency marginals. (d) Normalized lifetime-FRET histograms for increasing potential timescales (1 *µ*s, blue; 0.5 *µ*s, purple; 0.1 *µ*s, red) at fixed internal diffusivity *D* = 0.25 Å */*ns. Smaller mean-reversion rate (larger *τ*_*U*_ ) yields a stronger deviation from the static FRET line and a larger apparent shift in the mean FRET efficiency. (e) Lifetime-FRET histograms in the presence of an acceptor triplet state for two internal diffusivities (*D*_1_ = 0.1 Å^2^ */*ns, orange/green/purple; *D*_2_ = 1.0 Å^2^ */*ns, red/yellow/cyan) and three intersystem crossing quantum yields (*φ*_isc_ = 0; purple, blue; 1.1 *×*10^−3^; green, yellow; 2.5 *×*10^−3^; orange, red). Larger internal diffusivity reduces the apparent impact of the triplet state by enhancing conformational averaging. Inset: shift in mean FRET efficiency (circles) and per-photon, mean-subtracted divergence 𝒮 (squares) for species 1 (low *D*, purple/orange) and species 2 (high *D*, cyan/red). (f) Lifetime-FRET histograms for two species with different potential timescales (*τ*_*U*,1_ = 100 ns and *τ*_*U*,2_ = 500 ns) but fixed internal diffusivity. A smaller mean-reversion rate produces a larger decrease in mean FRET efficiency, analogous to panel (d), and renders the relative influence of the triplet state less prominent. Inset: shift (circles) and per-photon divergence 𝒮 (squares) of FRET histograms for species 1 (fast *τ*_*U*_, purple/orange) and species 2 (slow *τ*_*U*_, cyan/red).

Introducing an acceptor triplet state adds a second, photophysical route to low-FRET events, whose impact depends sensitively on the underlying continuum dynamics. For fixed (*τ*_*U*_, *D*), the triplet state simply introduces an additional effective “dark” contribution whose weight is set by the mean triplet fraction 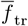, and the leadingorder mean shift remains approximately linear in both 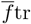 and the underlying FRET efficiency – as shown by the linear character of the circles of the insets in panels e-f and described in Sec. III. In Fig. 5e, this manifests as a larger shift and larger per-photon divergence 𝒮 (i.e., dynamic broadening) for the higher-diffusivity and higher-FRET populations, whereas lower apparent FRET efficiencies are visibly less affected, as highlighted in the insets. Increasing *D* enhances conformational averaging and partially masks the triplet signature, making the same intersystem crossing quantum yield appear less consequential for rapidly fluctuating chains than for more localized ones. A similar trend arises when varying *τ*_*U*_ at fixed *D*, Fig. 5f: slower potential relaxation produces a more substantial decrease in the mean FRET efficiency through conformational dynamics alone, while simultaneously reducing the relative contribution of the triplet-induced attenuation. In this sense, conformational entropy and slow reconfiguration can mimic, and in some regimes overshadow, the effects of triplet blinking on the observables.

## V. CONCLUSIONS

smFRET remains a cornerstone of experimental biophysics, owing to its ability to probe biomolecular structure and dynamics on nanometer and microsecond scales. At the same time, its reliance on fluorescent reporters necessarily entangles conformational observables with photophysical processes such as triplet blinking and photobleaching. To demonstrate the utility of a novel, computationally efficient framework for simulating confocal fluorescence microscopy – built around the concatenation of Brownian bridges – we carried out a quantitative analysis of how acceptor triplet states modulate the apparent conformational dynamics in PIE-based smFRET. Conformational motion was treated both in a discrete-state picture, with transitions between finitely many minima described by a CTMC, and in a continuum picture, with donor-acceptor separation obeying a modified OU process.

Across both representations, we identified clear dynamical regimes in which triplet blinking either averages out, produces conformation-specific attenuation, or overlaps nontrivially with conformational exchange, thereby reshaping FRET histograms and lifetime-based observables. In particular, we showed how intermediate-timescale overlap can partially homogenize FRET distributions and bias lifetime estimates in ways that may be misinterpreted as weaker or slower conformational dynamics if photophysics is not explicitly modeled.

The present framework suggests several natural extensions. Foremost among these is the explicit treatment of fluorophore and linker motions, including dye rotational diffusion and local configurational dynamics, which becomes tractable here because the simulation framework permits high temporal resolution while retaining numerical stability and computational efficiency. A potential approach to this problem, building on the recent works of Refs. [10, 27], is presented in Appendix A. More broadly, the Brownian-bridge-based photon simulation scheme is not limited to smFRET, but can be straightforwardly adapted to other confocal fluorescence techniques. In particular, extending the present methodology to fluorescence correlation spectroscopy (FCS) and related correlation-based measurements (e.g., FRET-FCS) would enable forward simulations of intensity-time traces with (sub)nanosecond resolution, including realistic photophysics and complex conformational dynamics within the confocal volume. The Julia code used to generate all simulations and analyses presented in this work is available as open source at https://github.com/LabGradinaru/SiMFRET.

## ACKNOWLEDGMENTS

We wish to thank Chifeng Wang for in aiding in the development of the tools for fluorescence lifetime estimation. The vast majority of simulations reported in this article were performed on the University of Toronto Mississauga High Performance Computing cluster and we thank Kenji Li for his help in configuring these resources. This work has been supported by the Natural Sciences and Engineering Research Council of Canada (NSERC RGPIN-2023-04864 to C.C.G.).

## Appendix A: Aliphatic linker and fluorophore motions

When we lift the assumption of transition dipole translational isotropy, instead of considering a single vector quantity ***R***, describing the donor-to-acceptor displacement, the individual contributions of the molecule and the linkers to the conformational dynamics must be considered,

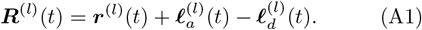

Here, ***ℓ***_*d*_ (resp. ***ℓ***_*a*_) describes the motion of aliphatic linker connecting donor (resp. acceptor) fluorophore to the molecule and ***r*** denotes the displacement between the molecule’s residues where the linkers anchor to the biomolecule. The superscript (*l*) indicates laboratory-frame quantities. In effect, two additional vector quantities for the linker motions must be added to the trajectory resultant from the conformational dynamics.

To model the rigidity of the molecule and linkers, four unit vectors are introduced to represent directions tangent to the polymer at the two linker attachment sites, 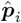 for *i* ∈*{d, a}*, and at the two *σ*-bonds connecting the last mobile linker backbone atom and the fluorophoreattachment atom, 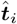, as shown in Fig. 6. Finally, to allow for additional freedom in the electronic transition dipole motion [28], unit vectors 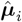 specifying their direction, are included in the model.

**FIG. 6.**
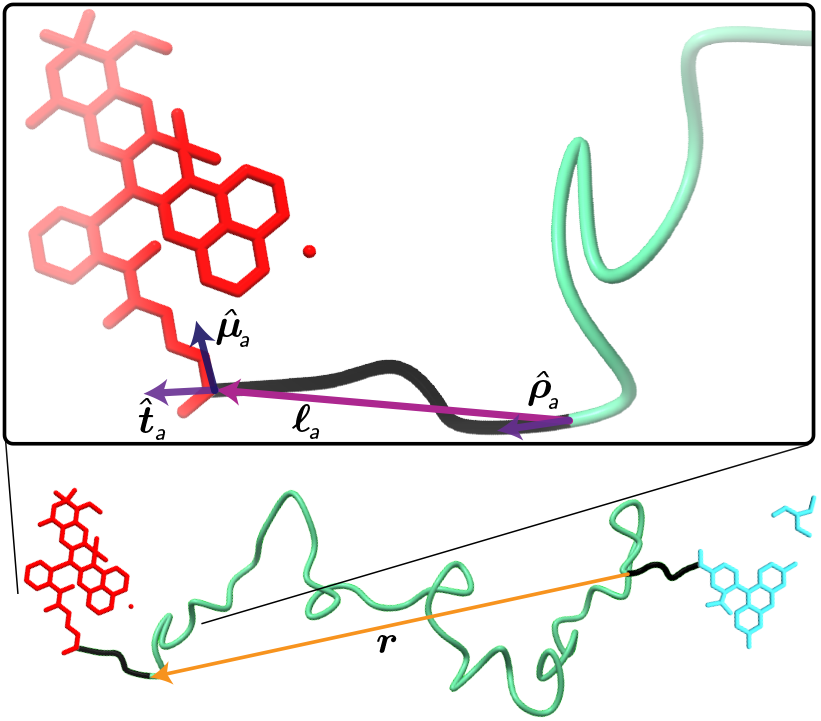
Instantaneous configuration of a sample biomolecule illustrating the proposed nine vector parameterization of molecule and linker dynamics in FRET.

It is often both more natural and computationally efficient to represent rigid-body rotations by unit quaternions (versors). A versor is a four-vector,

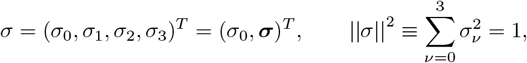

with *σ* and −*σ* encoding the same spatial rotation. The (non-abelian) group law is the Hamilton product: for *σ* = (*σ*_0_, ***σ***)^*T*^ and *ζ* = (*ζ*_0_, ***ζ***)^*T*^ [29],

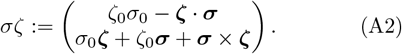

The inverse is the quaternion conjugate, *σ*^−1^ = *σ*^*^ := (*σ*_0_, *σ*_1_, *σ*_2_, *σ*_3_). If *σ* maps the body frame to the laboratory frame, a body-frame vector ***v***^(*b*)^ ∈ ℝ ^3^, understood in versor-based calculations as a pure quaternion embedding (0, ***v***^(*b*)^) ∈ ℚ, is rotated to the lab frame by the product ***v***^(*l*)^ = *σ****v***^(*b*)^*σ*^*^, which is equivalent to multiplying by the rotation matrix *R*(*σ*) constructed from *σ* [30].

As a heterogeneous, rotational analogue of the over-damped translational dynamics in Eq. (6), consider a torque ***τ*** ^(*l*)^ evaluated in the in the laboratory frame. Rotation to the body frame can be achieved by inversion of the above rule, ***τ*** ^(*b*)^ = *σ*^*^***τ*** ^(*l*)^*σ*. If the system has body-frame diffusivity 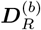, the overdamped rotational SDE for the body-to-lab versor can be written compactly as [31]

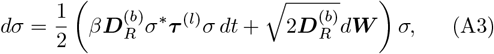

for inverse temperature *β* = (*k*_*B*_*T* )^−1^. Right-multiplication by *σ* translates the otherwise pure, body frame quaternions into genuine versors; numerically, one may enforce ||*σ*|| = 1 by a computationally inexpensive renormalization after each time step.

Centre of mass (COM) rotation of the biomolecule is modelled by a freely rotating sphere with fixed, effective rotational diffusivity, i.e., Eq. (A3) with ***τ*** ^(*l*)^ = 0. This coarse-grained description omits short-lived correlations between conformational fluctuations and overall rotation, yet it retains the dominant stochastic reorientation responsible for macroscopic depolarization and modulation of the end-to-end distance vector on the experimental timescale. Since our primary interest is the rotational motion local to the fluorophores – which is modelled with greater fidelity – any residual anisotropy in the global rotational diffusivity is expected to have only a minor effect, unless it is extreme.

Within the COM frame, for a highly disordered biomolecule whose contour length greatly exceeds its hydrodynamic radius, local motions fo tangent vectors to the molecule may be treated as statistically independent of the global COM rotation. Without loss of generality, let 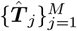 denote unit tangent vectors along the polymer or linker. Neighboring tangents are elastically coupled; a coarse-grained bending stiffness *K*_*j*−1_ penalizes curvature of the intervening worm-like chain (WLC) segment. A convenient representation of the deterministic restoring torque on the *j*th tangent, in the body frame, is

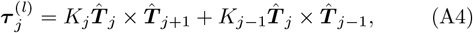

which is consistent with quadratic WLC bending penalties. In the WLC picture, the effective couplings scale as *K*_*j*_ ∝ *b*_*j*_*/L*_*j*_ where *b*_*j*_ is the local persistence length and *L*_*j*_ the contour length between nodes; thus *K*_*j*_ → 0, and tangent motions decorrelate, when *b*_*j*_ → 0 or *L*_*j*_ → ∞. At chain boundaries, the torque reduces to 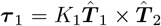 and 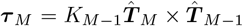. In the present application, we consider the case of four tangents, 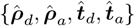, with the linker tangents 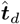 and 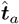 at the ends, see Fig. 6.

Let *ζ*_*j*_ be the versor mapping the *j*th tangent-fixed reference frame into the body frame. Assuming locally isotropic rotational diffusivity *D*_*R,j*_ in the tangent frame, the overdamped quaternion dynamics consistent with Eq. (A3) are

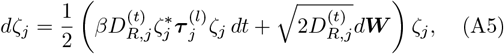

for torques 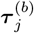 as stated in Eq. (A4). Eq. (A5) is the quaternion analogue of a Rouse model whereby springlike couplings coarse grain intramolecular interactions.

Up to this point, the motion of the linker itself has not been treated explicitly, beyond its coupling to the molecular coordinates in Eq. (A5). We distinguish two principal contributions to linker dynamics: (i) fluctuations of the effective contour length, arising from stretching/compression of the series of carbon-carbon bonds that compose the linker, and (ii) local steric interactions between the linker-fluorophore pair and the biomolecular surface. The former can be effectively modeled, as in the molecule itself in the main text, by an OU process for the end-to-end linker length, with mean-reversion rate *k* and equilibrium length *ℓ*_eq_.

Steric effects are more system-specific and generally depend on the detailed molecule topology and labeling geometry. In the present coarse-grained model, we assume that the fluorophore is bulkier than the linker and resides at its distal end, so that steric interactions can be taken to act at the tip of the linker vector, at distance *ℓ* = ||***ℓ***||. We approximate the local molecular surface as a right circular cone with opening angle 0 ≤ *ψ < π*, and describe the short-range repulsion by an inverse-powerlaw potential (e.g., Lennard-Jones) with exponent *α >* 0 and characteristic length scale *λ*. A simple geometric construction then yields the coupled angular-radial steric potential

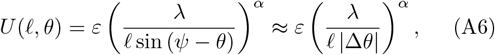

where *ℓ* is the end-to-end linker length, *θ* is the polar angle of the linker relative to the cone axis, Δ*θ* = *ψ* − *θ*, and *ε* sets the energy scale of the steric repulsion. The small-angle approximation emphasizes the rapid growth of steric cost as the dye approaches the conical boundary, providing a simple, tunable barrier that couples rotational and extensional degrees of freedom.

Including this steric term, the coupled radial-angular dynamics of the linker are governed by

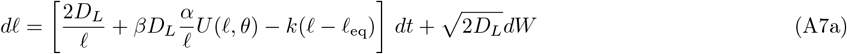

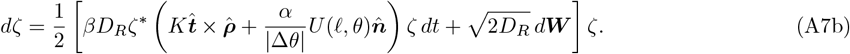

Here, *D*_*L*_ and *D*_*R*_ are the linker translational and rotational diffusion coefficients, respectively, *ζ* is the unit quaternion describing the fluorophore orientation, 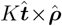 represents the intrinsic bending torque, see Eq. (A4), and 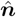 is the unit vector pointing along increasing polar angle in the center-of-mass frame. The latter can be written as

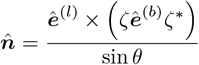

for fixed reference vectors ***ê***^(*l*)^ and ***ê***^(*b*)^ in the lab and body frames, respectively. Donor and acceptor linkers, and their associated parameters, are distinguished by subscripts *d, a* in Eqs. (A7a) and (A7b) as needed.

Qualitatively, as the dye approaches the molecular surface, either through linker compaction (*ℓ* → 0) or by grazing the conical boundary (Δ*θ* → 0), the steric term generates a large restoring force and torque that rapidly reorients and extends the linker away from the surface. Even at a heuristic level, this framework explains the breakdown of a pure OU description for linker-fluorophore orientation dynamics (cf. Ref. [27]): inhomogeneous angular sampling emerges naturally from the combined action of a WLC-like bending torque and a short-range steric barrier, leading to anisotropic orientational ensembles and non-Gaussian fluctuations in the effective dye-dye distance and orientation factor.

Using the machinery developed above, all direction-dependent factors are evaluated by rotating between frames with versors, using the rotational invariance of the dot product. Let *σ* map the body frame to the laboratory frame; then the lab polarization 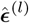 expressed in the body frame is 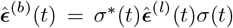. Similarly, for a linker or transition dipole specified in its tangent frame by ***û***^(*t*)^ and evolved by *ζ*(*t*) (cf. Eqs. (A5), (A7b)), 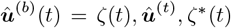. The donor-acceptor displacement in the body frame, consistent with Eq. (A1), is then

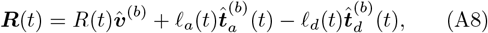

where 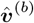 is a fixed body-frame unit vector describing the direction along which molecular distance fluctuations occur. All together, the time-dependent rates, *k*_fret_ and *k*_ex_ in Eqs. (1), (3), become explicit functionals of the stochastic coordinates.

