## Supplementary material for "Parsing contributions of physical phenomena to smFRET statistical inhomogeneity via multiparameter stochastic simulations": Suplemental Material

Aiyan Brown and Claudiu C. Gradinaru
*Department of Physics, University of Toronto, 60 Saint George St., Toronto, Ontario, M5S 1A7, Canada and Department of Chemical and Physical Sciences, University of Toronto Mississauga, 3359 Mississauga Road, Mississauga, Ontario, L5L 1C6, Canada*
(Dated: December 5, 2025)

### I. CHOICE OF EFFECTIVE RADIUS

It is well known (cf. [1, 2]) that the mean first passage time (MFPT) $\mathcal{M}(\mathbf{x})$ of a particle with unit diffusivity from a unit sphere in $d$-dimensions satisfies a Poisson equation,

$$\nabla^2\mathcal{M}(\mathbf{x}) = -1. \tag{S1}$$

A spherically-symmetric solution in $d$-dimensions can be read off simply as [1]

$$\mathcal{M}(\mathbf{x}) = \frac{1-||\mathbf{x}||^2}{2d}. \tag{S2}$$

In the case where instead we are interested in the exit time from an ellipsoid with semi-axes $a_i$, we can rescale coordinates as $x_i \to z_i = x_i/a_i$, mapping the ellipsoid to a sphere. We proceed with the ansatz that, in these new coordinates, the transformed MFPT, $\mathcal{Q}$, has the same quadratic form,

$$\nabla^2\mathcal{M} = \sum_i \frac{1}{a_i^2}\frac{\partial^2\mathcal{Q}}{\partial z_i^2} = -2\alpha\sum_i a_i^{-2} = -1, \tag{S3}$$

for a constant $\alpha$, which gives an explicit solution:

$$\mathcal{Q}(\mathbf{z}) = \frac{1-||\mathbf{z}||^2}{2\sum_i a_i^{-2}}. \tag{S4}$$

We understand the effective radius as that which translates the MFPT for a circle, i.e. $\mathcal{Q}(\mathbf{x}) = R_{\text{eff}}^2\mathcal{M}(\mathbf{x})$, so that, for an ellipse,

$$R_{\text{eff}}^2 = \frac{d}{\sum_i a_i^{-2}}, \tag{S5}$$

where we have used dimensional analysis to deduce the scaling of $\mathcal{M}$ relative to $\mathcal{Q}$.

### II. TRIPLET-INDUCED SHIFT IN MEAN FRET EFFICIENCY

In the main text, we claimed that for an input FRET efficiency

$$\varepsilon = \frac{\hat{n}_a}{\hat{n}_a+\hat{n}_d},$$

the expected shift in the mean FRET efficiency due to an acceptor triplet state with time-averaged occupation fraction $\overline{f}$tr is linear in both $\varepsilon$ and $\overline{f}$tr. Here we outline a simple probabilistic argument supporting this statement.

Suppose the probability that an excited donor fluorophore deexcites radiatively (i.e., emits in the donor channel) in the absence of an acceptor triplet is

$$\mathbb{P}_{d_{\text{em}}|\text{no triplet}} = (1-\varphi_{\text{ic}})(1-\varepsilon),$$

where $\varphi_{\text{ic}}$ is the internal conversion quantum yield from the donor excited singlet to the ground state. Correspondingly, the acceptor emission probability (via FRET) is

$$\mathbb{P}_{d_{\text{em}}|\text{no triplet}} = (1-\varphi_{\text{ic}})\varepsilon$$

When the acceptor is in the triplet state, FRET is blocked and the donor must deexcite either radiatively or via internal conversion. If the acceptor spends a time-averaged fraction $\overline{f}_{\text{tr}}$ in the triplet state, then the donor emission probability becomes

$$\mathbb{P}_{d_{\text{em}}|\text{triplet}} = (1-\varphi_{\text{ic}})\left[\overline{f}_{\text{tr}} + (1-\overline{f}_{\text{tr}})(1-\varepsilon)\right],$$

while the acceptor emission probability is reduced to

$$\mathbb{P}_{a_{\text{em}}|\text{triplet}} = (1-\varphi_{\text{ic}})(1-\overline{f}_{\text{tr}})\varepsilon.$$

Thus the donor and acceptor channels are rescaled by factors

$$\alpha_d = \frac{\mathbb{P}_{d_{\text{em}}|\text{triplet}}}{\mathbb{P}_{d_{\text{em}}|\text{no triplet}}} = 1 + \frac{\overline{f}_{\text{tr}}\varepsilon}{1-\varepsilon}, \tag{S6a}$$

$$\alpha_a = \frac{\mathbb{P}_{a_{\text{em}}|\text{triplet}}}{\mathbb{P}_{a_{\text{em}}|\text{no triplet}}} = 1 - \overline{f}_{\text{tr}}. \tag{S6b}$$

Starting from burst-averaged counts $(\hat{n}_a, \hat{n}_d)$ in the absence of triplet, the corresponding counts in the presence of triplet blinking are approximately $\hat{n}'_a = \alpha_a\hat{n}_a$ and $\hat{n}'_d = \alpha_d\hat{n}_d$, amounting to an apparent FRET efficiency

$$\varepsilon' = \frac{\hat{n}'_a}{\hat{n}'_a+\hat{n}'_d}.$$

The shift in mean FRET efficiency is then

$$\Delta = \frac{\frac{\mathbb{P}_{a_{\text{em}}|\text{triplet}}}{\mathbb{P}_{a_{\text{em}}|\text{no triplet}}}\hat{n}_a}{\frac{\mathbb{P}_{a_{\text{em}}|\text{triplet}}}{\mathbb{P}_{a_{\text{em}}|\text{no triplet}}}\hat{n}_a + \frac{\mathbb{P}_{d_{\text{em}}|\text{triplet}}}{\mathbb{P}_{d_{\text{em}}|\text{no triplet}}}\hat{n}_d} - \varepsilon \tag{S7}$$

$$= \frac{(1-\overline{f}_{\text{tr}})\hat{n}_a}{(1-\overline{f}_{\text{tr}})\hat{n}_a + (1-\overline{f}_{\text{tr}} + \frac{\overline{f}_{\text{tr}}}{1-\varepsilon})\hat{n}_d} - \frac{\hat{n}_a}{\hat{n}_a - \hat{n}_d} \tag{S8}$$

$$= -\overline{f}_{\text{tr}}\varepsilon. \tag{S9}$$

For reference, the projections of Fig. 3c in the main text – by plotting the absolute shift in mean FRET efficiency as a function of input FRET efficiency and time-averaged triplet fraction – are shown in Fig. S1.

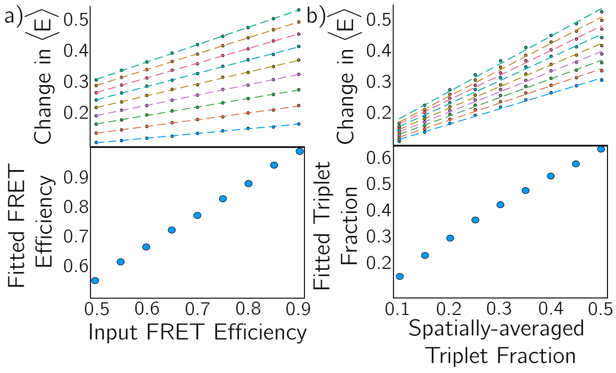


FIG. S1. Change in mean FRET efficiency as a function of input FRET efficiency (a; top) and triplet fraction (b; top) for FRET efficiencies $\{0.5, 0.55, \ldots, 0.9\}$ and triplet fractions $\{0.1, 0.15, \ldots, 0.5\}$. The linear relationship between each of the coordinates is demonstrated by performing an unweighted linear regression, $\Delta(x) = ax + b$ for each set of parameters; each of which returned a coefficient of determination of at least 0.995 and $|b| < 0.065$. (bottom) Overestimate of fitted results, based on fitting to $-\overline{f}_{\rm tr}\varepsilon$, for both the triplet fraction and apparent FRET efficiency.

The simulated relationships between $\Delta$ and each of $\overline{f}_{\rm tr}$ and $\varepsilon$ are clearly linear, consistent with Eq. (S9), but the fitted slopes exhibit small systematic deviations from the ideal value of $+1$. We attribute these discrepancies to be a consequence of the triplet fraction used in the analysis being computed from a spatio-temporally averaged excitation rate, which is only an approximation to the true excitation kinetics. In particular, for pulsed excitation the instantaneous peak intensity within the confocal volume is higher than suggested by the mean excitation rate, leading to a systematically higher true triplet occupation than that inferred from the averaged rate. As a result, the effective $\overline{f}_{\rm tr}$ entering Eq. (S9) is underestimated, and the apparent slope of $\Delta$ versus $\overline{f}_{\rm tr}\varepsilon$ is correspondingly reduced. Within this approximation, however, the leading-order dependence $\Delta \propto -\overline{f}_{\rm tr}\varepsilon$ still provides an accurate and useful description over the parameter ranges considered here.

In the main text, "heatmaps" were used in order to more effectively display the data. For completeness, the set of lifetime-FRET plots used to construct these are included in Fig. S2. Qualitatively, and consistent with other choices of input FRET efficiency, the maximal (mean-subtracted) divergence occurs at the triplet fractions for which the "dynamic shift" [5] is largest along the corresponding dynamic FRET line.

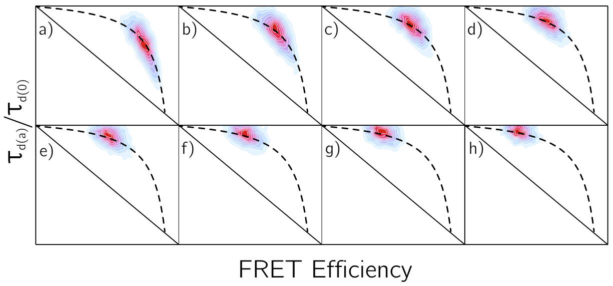


FIG. S2. Two-dimensional histograms for fixed, $\varepsilon = 0.9$, input FRET efficiency with varying triplet fraction; (a) $\overline{f}_{\rm tr} = 0.10$, (b) $\overline{f}_{\rm tr} = 0.15$, ..., (h) $\overline{f}_{\rm tr} = 0.50$. The dashed line is the dynamic FRET line $s(E) = \hat{\tau}^{(1)}_{d(a)} + \hat{\tau}^{(2)}_{d(a)} - \hat{\tau}^{(1)}_{d(a)}\hat{\tau}^{(2)}_{d(a)}/(1-E)$ [3, 4], where $\hat{\tau}^{(i)}_{d(a)}$ is the normalized lifetime of the $i$th population (here, $\hat{\tau}^{(1)}_{d(a)} = 1.0$ and $\hat{\tau}^{(2)}_{d(a)} = 0.1$).

### III. PER-PHOTON DIVERGENCE AND THE SHOT-NOISE LIMIT

We recall that the Kullback-Leibler (KL) divergence between two distributions with probability mass functions $p$ and $q$ on a discrete set $\Omega$ is

$$D_{\rm KL}(p \,||\, q) = \sum_{x\in\Omega} p(x) \log \frac{p(x)}{q(x)}. \qquad \text{(S10)}$$

Here, we take $q$ to be the shot-noise limit (SNL) of an smFRET histogram, i.e., the distribution that would be obtained from an infinitely long (ergodic) measurement of a sample with a single underlying FRET efficiency. This reference distribution was derived by Nir *et al.* [6] as

$$q_\varepsilon(x) = \frac{1}{n_{\rm bursts}} \sum_{n=0}^{\infty} f(n)\mathbb{P}_\varepsilon(nx \mid n)\mathbf{1}_{\mathbb{N}}(nx), \quad x \in \mathbb{Q}\cap[0,1], \qquad \text{(S11)}$$

where $\varepsilon$ is the true (input) FRET efficiency, $x$ is the FRET estimator (the "proximity ratio"), $f(n)$ is the probability mass function of burst sizes (in photon counts), $\mathbb{P}_\varepsilon(\cdot \mid n)$ is the conditional distribution of the number of acceptor photons given $n$ total photons, taken to be binomial, and $\mathbf{1}_A(x)$ is the indicator function – one if $x \in A$ and zero otherwise.

In practice, an empirical estimate of $q_\varepsilon$ can be constructed directly from experimental data. Given a measured burst-size distribution $f(n)$, one samples burst sizes $n$ from $f$, then for each $n$ draws multiple realizations from the binomial distribution $\mathbb{P}_\varepsilon(\cdot \mid n)$, converts the resulting acceptor counts to proximity ratios $x$, and accumulates a histogram over $x$. Under the assumptions of ergodicity and that the measured $f(n)$ faithfully reflects the true burst-size distribution, repeated sampling yields a Monte Carlo approximation that converges to the SNL $q_\varepsilon$ as the number of realizations increases. The divergence can be subsequently calculated by partitioning the domain on which the FRET efficiency is defined,

$[0, 1]$, and evaluating Eq. (S10) on all partitions on which $p$ is non-zero.

We believe that the mean-subtracted, per-photon KL divergence $\mathcal{S}$ is a more robust estimator of dynamics than the frequently used comparison of the standard deviations of the SNL and observed distributions – an approach that underlies, for example, burst-variance analysis [7] and its time-resolved variants [8]. First, $D_{\mathrm{KL}}$ is sensitive to the full shape of the distribution (including skewness and tail behaviour), rather than compressing all deviations into a single second moment. Second, by subtracting the KL divergence of an SNL distribution with the same mean, we largely factor out static heterogeneity and mean shifts (e.g., due to triplet-induced attenuation), isolating the genuinely dynamical contribution to the broadening. Finally, the KL divergence is dimensionless and naturally comparable across different FRET efficiencies and burst-size distributions, making it well suited for quantitative mapping of dynamical regimes.

To illustrate the behaviour of $\mathcal{S}$, consider a molecule whose conformational dynamics can be modelled by two discrete states with FRET efficiencies $E_1$ and $E_2$, as shown in Fig. S3. Let $0 \leq \phi \leq 1$ denote the steady-state occupancy fraction of state $E_1$, and let $\chi$ be a burst-wise estimator of $\phi$ (so that $\mathbb{E}[\chi] = \phi$). For a given burst, the probability of detecting an acceptor photon upon donor excitation is

$$P_a(\chi) = \chi E_1 + (1-\chi)E_2 = E_2 + \chi(E_1 - E_2). \quad \text{(S12)}$$

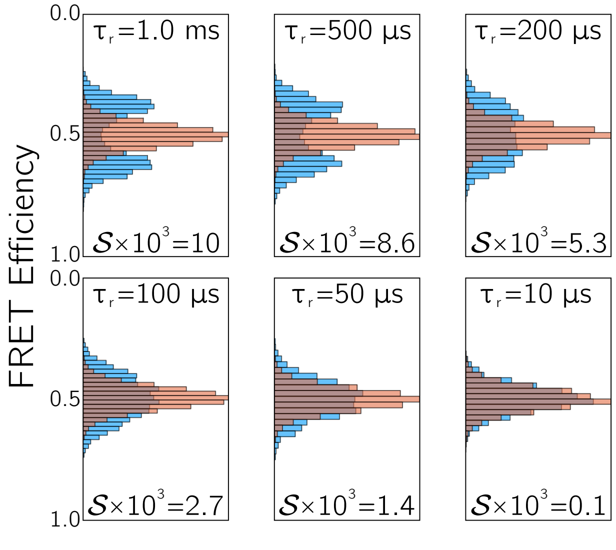


FIG. S3. Comparison between the shot-noise limit (SNL; orange) and the simulated FRET distribution (blue) for a molecule undergoing two-state conformational exchange, shown for several interconversion times $\tau_r$. As $\tau_r$ decreases and conformational exchange becomes fast on the burst timescale, the simulated distributions converge toward the SNL, reflecting the effective averaging over $E_1$ and $E_2$. The underlying FRET efficiencies of the two conformational states are kept fixed at $E_1 = 0.35$ and $E_2 = 0.65$ in all cases.

Conditional on $\chi$ and a burst containing $N$ photons, the acceptor counts are taken to be shot-noise limited, $n_a \mid \chi \sim \text{Binomial}\,(N, P_a(\chi))$. Defining the burst-wise FRET estimator $\varepsilon = n_a/N$, the law of total expectation gives

$$\mathbb{E}[\varepsilon] = \mathbb{E}\chi\,[\mathbb{E}[\varepsilon \mid \chi]] = \mathbb{E}\chi\,[P_a(\chi)] = \phi E_1 + (1-\phi)E_2, \quad \text{(S13)}$$

which recovers the population-weighted mean efficiency, as expected.

Similarly, for the variance, by application of the law of total variance, since $\varepsilon$ is Bernoulli distributed given $\chi$,

$$\mathrm{Var}\,[\varepsilon] = \mathbb{E}_\chi\,[\mathrm{Var}\,[\varepsilon \mid \chi]] + \mathrm{Var}\,[\mathbb{E}[\varepsilon \mid \chi]] \quad \text{(S14)}$$

$$= \underbrace{\frac{1}{N}\mathbb{E}_\chi\left[P_a(\chi)(1 - P_a(\chi))\right]}_{\sigma_0^2} + \underbrace{\mathrm{Var}\left[(P_a(\chi))\right]}_{\sigma_{\mathrm{dyn}}^2} \quad \text{(S15)}$$

where the first term, $\sigma_0^2$, is the shot-noise contribution (the SNL variance for the given burst-size distribution), and the second term,

$$\sigma_{\mathrm{dyn}}^2 = \mathrm{Var}\,[P_a(\chi)] = (E_1 - E_2)^2 \mathrm{Var}\,[\chi]\,, \quad \text{(S16)}$$

is the dynamical contribution arising from fluctuations of the occupancy fraction. The quantity $\mathrm{Var}\,[\chi]$ implicitly encodes the dynamical timescale: if the interconversion time $\tau_r$ is large compared to the burst duration, the slow-exchange limit, $\chi$ becomes approximately Bernoulli-distributed with parameter $\phi$, giving $\mathrm{Var}\,[\chi] \approx \phi(1-\phi)$ and a maximal dynamical contribution – see, for example, Fig. 4b of the main text. To extend this analysis to smaller $\tau_r$, let $n = \lfloor T/\tau_r \rfloor$ denote the effective number of statistically independent "exchange events" within a burst of length $T$. Approximating the trajectory as piecewise constant over these blocks, we can write

$$\chi = \frac{1}{n}\sum_{k=1}^{n} \mathbf{1}_k, \quad \text{(S17)}$$

where $\mathbf{1}_k$ is an indicator variable taking value 1 if the system is in state $E_1$ during block $k$ and 0 otherwise. In the simplest approximation, the $\mathbf{1}_k$ are i.i.d. Bernoulli($\phi$), so

$$\mathrm{Var}\,[\chi] = \frac{1}{n^2}\sum_{k=1}^{n} \mathrm{Var}\,[\mathbf{1}_k] = \frac{\phi(1-\phi)}{n} \approx \phi(1-\phi)\frac{\tau_r}{T}. \quad \text{(S18)}$$

This expression interpolates smoothly between the slow-exchange limit ($\tau_r \gg T$, effectively $n \to 1$, $\mathrm{Var}\,[\chi] \to \phi(1-\phi)$) and the fast-exchange limit ($\tau_r \ll T$, $n \gg 1$, $\mathrm{Var}\,[\chi] \to 0$). Substituting into Eq. (S16), we obtain the approximate scaling

$$\sigma_{\mathrm{dyn}}^2 \approx (E_1 - E_2)^2 \phi(1-\phi)\frac{\tau_r}{T}, \quad \text{(S19)}$$

which achieves a maxima at $\phi = 0.5$.

One can extend this analysis to higher-order moments, but even this second-moment decomposition is informative. If we (crudely) approximate both the SNL and observed FRET distributions as Gaussian with the same mean and variances $\sigma_0^2$ and $\sigma_0^2+\sigma_{\rm dyn}^2$, respectively, $\mathcal{S}$ can be evaluated in closed form as

$$\mathcal{S} = \frac{1}{2\langle N\rangle}\left[\frac{\sigma_{\rm dyn}^2}{\sigma_0^2} - \log\left(1+\frac{\sigma_{\rm dyn}^2}{\sigma_0^2}\right)\right], \quad \text{(S20)}$$

where $\langle N\rangle$ is the average of the burst size distribution. This compensates for the scaling $\sigma_0^2 \sim N^2$ of the SNL variance [6]. In the specific case of acceptor triplet dynamics discussed in the main text, an apparent state with null FRET efficiency is induced, so we may take $E_1 = 0$ and $E_2 = \varepsilon$ (the underlying non-zero efficiency). Then $\sigma_{\rm dyn}^2 \propto \varepsilon^2 {\rm Var}[\chi]$, and for fixed dynamical amplitude ${\rm Var}[\chi]$, $\mathcal{S}$ increases monotonically with the input FRET efficiency. This is consistent with the behaviour observed in Fig. 3c of the main text and highlights how $\mathcal{S}$ encodes both the magnitude of conformational heterogeneity (via $E_1 - E_2$ and ${\rm Var}[\chi]$) and its interplay with the shot-noise level at a given $\varepsilon$. Notably, however, $\mathcal{S}$ varies non-trivially with triplet fraction, see Fig. S2, at odds with the scaling relation Eq. (S19), suggesting that higher-order moments contribute non-negligibly to $\mathcal{S}$ when the number of exchanges is large, shown qualitatively in Fig. S5.

### IV. BURST SIZE DISTRIBUTIONS

Burst-size distributions have a pronounced influence on the SNL and thus on the observed FRET histograms themselves. In experiment, there is an intrinsic tradeoff between excitation power and photobleaching propensity: increasing the excitation intensity boosts the detected photon rate and narrows the SNL, but also accelerates irreversible photo-destruction of the fluorophores. Typical time-averaged excitation powers in confocal smFRET setups lie in the range of 50-100 $\mu$W at the sample [9]. In addition, the overall detection efficiency of the microscope-detector system is far from unity, being limited by the finite numerical aperture of the objective, imperfect transmission through the optical train, detector quantum efficiency, and timing electronics. In practice, the net detection efficiency often lies near $\sim 10\%$.

In our simulations, these experimental limitations were mimicked by Bernoulli thinning of the detected photon streams: after photon generation, each photon was retained with a fixed probability and otherwise discarded. For the results presented in the main text, a 95% rejection probability (i.e., an effective detection probability of 5%) was used for all channels. This choice compensates both for a net detection efficiency of order 15% and for the approximate one-third reduction in apparent excitation rate associated with the pulsed, interleaved excitation scheme (PIE). In a strictly weak-excitation regime, reducing the excitation power or thinning the detected photons are equivalent up to an overall rescaling of the photon rate, so this treatment provides a convenient and computationally simple surrogate for the integrating the quaternion models discussed in the Appendix of the main text. We furthermore restricted ourselves to excitation powers comparable to typical experimental values – 150 $\mu$W of 532 nm "donor" excitation and 105 $\mu$W of 633 nm "acceptor" excitation – with fluorescence quantum yields of 0.5 and absorption cross sections $\sigma_{d,\rm abs} = \sigma_{a,\rm abs} = 10^{-20}$ m$^2$. Under these conditions, the average burst size was maintained near $\sim 200$ detected photons, split amongst the three PIE photon streams, as summarized in Fig. S4.

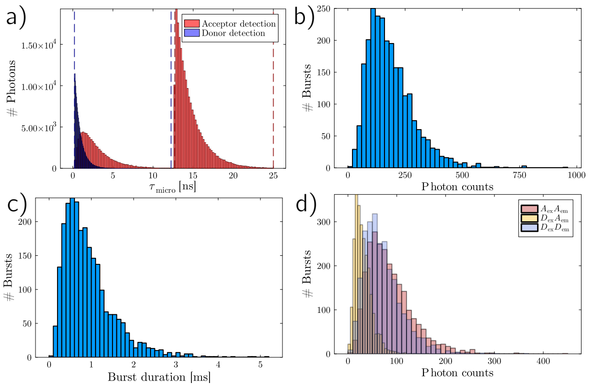


FIG. S4. Representative burst statistics used in the simulations. (a) Microtime histogram demonstrating the exponential donor-pulse excitation-emission cycle (blue, left), the bi-exponential FRET fluorescence decay (red, left) and exponential acceptor fluorescence decay (red, right). (b) Burst length distribution, measured as the time from the first to last photon in each burst, with a pinhole diameter $\varnothing = 50$ $\mu$m. (c) Burst size distributions, in numbers of photons, for all channels, with a 95% rejection probability. (d) Per-channel burst size distributions.

Given no burst search was performed, and rather the sum of all photons from a given molecule were collected to constitute a burst, the burst duration is longer than would be observed experimentally (Fig. S4c). Moreover, from our own measurements, an average of $\sim 200$ detected photons per burst is toward the upper end of what is routinely obtained for freely diffusing single molecules under PIE excitation. To assess the robustness of our conclusions with respect to the burst-size distribution, we therefore repeated the analysis underlying Fig. 3c of the main text while systematically varying the mean burst size. Operationally, this was achieved by changing the strength of the Bernoulli filter applied to the simulated photons, thereby tuning the effective detection probability and hence the mean number of detected photons per burst. The resulting dependence of the standard deviation and per-photon divergence $\mathcal{S}$ on input FRET efficiency and triplet fraction for several different target burst sizes is shown in Fig. S5.

These results highlight three key points. First, the mean FRET efficiency is largely insensitive to the burst-size distribution over the experimentally relevant range of photon numbers, consistent with the fact that the underlying population-weighted efficiency does not depend on detection efficiency. Second, the absolute standard deviation of the FRET histogram depends strongly on the

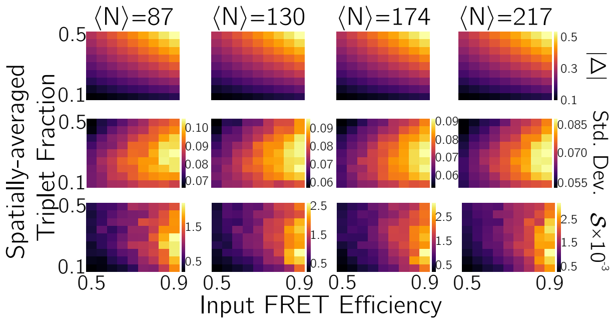


FIG. S5. Effect of burst size on acceptor triplet-induced FRET distortions. Top row: absolute shift in mean FRET efficiency as a function of input FRET efficiency and mean triplet fraction for three different mean burst sizes (increasing left to right). Middle row: standard deviation of the simulated FRET distributions under the same conditions. Bottom row: per-photon mean-subtracted KL divergence $\mathcal{S}$ between the SNL and the simulated FRET distributions.

mean burst size: dimmer/smaller bursts are dominated by shot noise and show broader distributions – increasing the upper and lower bound on the range of standard deviations – whereas brighter/larger bursts approach the asymptotic SNL and exhibit narrower histograms. Third, the per-photon divergence $\mathcal{S}$ is less sensitive to burst size than the raw variance and retains the same qualitative dependence on input FRET efficiency and triplet fraction. In particular, the loci of maximal divergence and the relative contrast between different dynamical regimes are essentially preserved across the tested photon-number conditions. This supports the interpretation of $\mathcal{S}$ as a more robust measure of dynamical broadening, largely factorized from trivial changes in photon statistics.
